# Genomic insights into the population history and biological adaptation of Southwestern Chinese Hmong-Mien people

**DOI:** 10.1101/2021.10.16.463767

**Authors:** Yan Liu, Jie Xie, Mengge Wang, Changhui Liu, Jingrong Zhu, Xing Zou, Wenshan Li, Lin Wang, Cuo Leng, Quyi Xu, Hui-Yuan Yeh, Chuan-Chao Wang, Xiaohong Wen, Chao Liu, Guanglin He

**Author notes:** These authors contributed equally to this work and should be considered co-first authors (Yan Liu, Jie Xie, Mengge Wang, Guanglin He). **Corresponding author** (**M.G. W.**), (**H.Y. Y.**), (**C.C. W.**), (**X.H. W.**), (**C. L.**) and (**G.L. H.**).

## Abstract

Hmong-Mien-speaking (HM) populations, widely distributed in South China, North of Thailand, Laos and Vietnam, have experienced different settlement environments, dietary habits and pathogen exposure. However, their specific biological adaptation also remained largely uncharacterized, which is important in the population evolutionary genetics and Trans-Omics for regional Precision Medicine. Besides, the origin and genetic diversity of HM people and their phylogenetic relationship with surrounding modern and ancient populations are unknown. Here, we reported genome-wide SNPs in 52 representative Miao people and combined them with 144 HM people from 13 geographically representative populations to characterize the full genetic admixture and adaptive landscape of HM speakers. We found that obvious genetic substructures existed in geographically different HM populations and also identified one new ancestral lineage specifically exited in HM people, which spatially distributed from Sichuan and Guizhou in the North to Thailand in the South and temporally dated to at least 500 years. The sharing patterns of the newly-identified homogeneous ancestry component combined the estimated admixture times via the decay of Linkage Disequilibrium and haplotype sharing in GLOBETROTTER suggested that the modern HM-speaking populations originated from Southwest China and migrated southward recently, which is consistent with the reconstructed phenomena of linguistic and archeological documents. Additionally, we identified specific adaptive signatures associated with several important human nervous system biological functions. Our pilot work emphasized the importance of anthropologically-informed sampling and deeply genetic structure reconstruction via whole-genome sequencing in the next step in the deep Chinese population genomic diversity project (CPGDP), especially in the ethnolinguistic regions.

## Introduction

Yungui Plateau and surrounding regions are the most ethnolinguistically diverse regions of China with a population size of approximately 205 million (2020 census), home to many ethnic groups, including the major population of Han Chinese and minorities of HM (HM), Tai-Kadai (TK), and TB (TB). This region is a mountainous and rugged area, consisting of Sichuan, Chongqing, Guizhou, Yunnan and most parts of Tibet Autonomous Region, which is characterized by the Sichuan Basin in the northeast, the karstic Yunnan-Guizhou Plateau in the east, and the Hengduan Mountains in the west and the majority of the region is drained by the Yangtze River. Historical records documented that portions of Southwest China were incorporated as unequivocal parts of greater China since at least the end of the third century BCE (Herman, 2018), and this region was largely dominated and incorporated into the Chinese domain by the time of the Ming dynasty (Harper, 2007). It has been suggested that the Nanman tribes were ancient indigenous people who inhabited in inland South and Southwest China. The Nanman referred to various ethnic groups and were probably the ancestors of some present-day HM, TK, and non-Sinitic Sino-Tibetan (ST) groups living in Southwest China. Generally, Southwest China exhibits a unique panorama of geographic, cultural, ethnic, linguistic, and genetic diversity. However, the complete picture of genetic diversity of ethnolinguistically divest populations in this region remained uncharacterized.

During the past decade, paleogenomic studies have transformed our knowledge of the population history of East Asians (Fu et al., 2013; Liu, Yichen et al., 2021; Mao et al., 2021; Ning et al., 2020; Ning et al., 2019; Wang, C.C. et al., 2021; Wang, T. et al., 2021; Yang et al., 2020). A recent archaeological study of early Holocene human cranium from Guizhou (Zhaoguo M1) supported that regionalization of morphological variability patterns between Neolithic northern and southern East Asians could trace back to at least 10,000 years ago (ya) (Zhang et al., 2021). However, our knowledge about the demographic history of populations in Southwest China is limited due to the lack of ancient DNA data and sparse sampling of modern people in genome-wide SNP or whole-genome studies (Bin et al., 2021; Chen et al., 2021b; Liu, Y. et al., 2021a; Wang, M. et al., 2021b; Wang et al., 2020). A series of recent genome-wide SNP studies demonstrated that southwestern Han Chinese showed a closer affinity with northern East Asian sources relative to indigenous populations and were well fitted via the admixture of ancient millet farmers from the Yellow River basin (YRB) and rice farmers from the Yangtze River basin (He et al., 2020; Liu, Y. et al., 2021a; Wang, M. et al., 2021a; Wang, M. et al., 2021b; Wang et al., 2020). Genetic findings focused on the culturally unique Hui people in this region also have proved that cultural diffusion has played an important role in the formation of the Hui people and southwestern Huis could be modeled as a mixture of major East Asian ancestry and minor western Eurasian ancestry (Liu, Y. et al., 2021a; Wang et al., 2020). He et al. further obtained genomic information from 131 TB-speaking Tujia individuals from Southwest/South-Central China and found the strong genetic assimilation between Tujia people and central Han Chinese, which provided evidence that massive population movements and genetic admixture under language borrowing have facilitated the formation of the genetic structure of Tujia people (He et al., 2020). The patterns of population structure of TK groups revealed the genetic differentiation among TK people from Southwest China and showed that YRB millet farmers and Yangtze River rice farmers contributed substantially to the gene pool of present-day inland TK people(Bin et al., 2021; Wang, M. et al., 2021a). Chen et al. recently analyzed genome-wide SNP data of 26 Mongolic-speaking Mongolians and 55 Tungusic-speaking Manchus from Guizhou and found that southwestern Mongolic/Tungusic groups had a stronger genetic affinity with southern East Asians than with northern Altaic groups (Chen et al., 2021b). It is remarkable, however, that no specific genome-wide studies have been published to shed new light on the population structure of HM groups from Southwest China.

Currently, HM groups mainly dwell in South China (including South-Central, Southwest, and Southeast China) (He et al., 2019; Huang et al., 2020; Xia et al., 2019; Zhang et al., 2019) and Vietnam, Laos and Thailand in mainland Southeast Asia (Kutanan et al., 2021; Liu et al., 2020). The history of the HM language family is obscure, which has been passed down mainly through oral legends and myths, for few written historical records exist. Hence, linguistic, genetic and paleogenomic studies are crucial for reconstructing the demographic history of HM groups (Huang et al., 2020; Kutanan et al., 2021; Liu et al., 2020; Wang, T. et al., 2021; Xia et al., 2019). Wang et al. successfully obtained genomic material from 31 ancient individuals from southern China (Guangxi and Fujian) ranging from ~12,000-10,000 to 500 ya and identified HM-related ancestry represented by ~500-year-old GaoHuaHua population (Wang, T. et al., 2021). Neolithic genomes from Southeast Asia also identified at least five waves of southward migrations from China participating in the formation of their modern patterns of genetic and ethnolinguistic diversity (Lipson et al., 2018; McColl et al., 2018). The genetic information of HM groups from South-Central China showed that HM-related ancestry was phylogenetically closer to the ancestry of Neolithic mainland Southeast Asians and modern Austroasiatic (AA) groups than to Austronesians (Xia et al., 2019). Huang et al. analyzed genome-wide SNP data of HM groups from Guangxi (Southeast China) and found that HM-related ancestry maximized in the western Hmong groups (Miao_Longlin and Miao_Xilin) (Huang et al., 2020). Findings of the human genetic history of mainland Southeast Asia also confirmed the observed heterogeneity in HM people derived from multiple ancestral sources during the extensive population movements and interactions (Kutanan et al., 2021; Liu et al., 2020). Therefore, systematic genome-wide studies focusing on southwestern Chinese HM groups and publicly available ancient East Asians will provide additional insights into the genetic makeup of HM groups from South China.

Here we generated new genome-wide data of 52 the northernmost HM-speaking Miao individuals from Xuyong, Sichuan. The Miao people are the largest of the HM-speaking populations and the fourth largest of the 55 ethnic minorities in China. The Miaos are a group of linguistically-related people mainly living in mountainous areas of South China. The Xuyong is a county in the southeastern of Sichuan province, which borders Guizhou to the south and Yunnan to the west. To thoroughly investigate the demographic history of southwestern HM groups, we co-analyzed newly generated data with publicly available genome-wide data of present-day and ancient East Eurasians leveraging shared alleles and haplotypes.

## Methods and materials

### Sample collection, genotyping and data merging

All newly-genotyped individuals were collected from three geographically different populations in Sichuan (Baila (14), Hele (19) and Jiancao (19), **Figure S1**). Oragene DN salivary collection tube was used to collect salivary samples. This study was approved via the ethical board of North Sichuan Medical College and followed the rules of the Helsinki Declaration. Informed consent was obtained from each participated volunteer. To keep a high representative of our included samples, the included subjects should be indigenous people and lived in the sample collection place for at least three generations. We genotyped 717,227 SNPs using the Infinium R Global Screening Array (GSA) version 2 in Miao people, which included 661,133 autosomal SNPs and the remaining 56,096 SNPs localized in X-/Y-chromosome and Mitochondrial DNA. We used PLINK (version v1.90) (Chang et al., 2015) to filter-out raw SNP data based on the missing rate (mind: 0.01 and geno: 0.01), allele frequency (--maf 0.01) and p values of Hardy-Weinberg exact test (--hwe 10^−6^). We used the King software to estimate the degrees of kinship among 52 individuals and remove the close relatives within the three generations (Tinker and Mather, 1993). We finally merged our data with publicly available modern and ancient reference data from Allen Ancient DNA Resource (AADR, https://reich.hms.harvard.edu/allen-ancient-dna-resource-aadr-downloadable-genotypes-present-day-and-ancient-dna-data), Besides, we also merged our new dataset with modern population data from China and Southeast Asia, and ancient population data from Guangxi, Fujian and other regions of East Asia (Mao et al., 2021; Wang, C.C. et al., 2021; Wang, T. et al., 2021; Yang et al., 2020) and finally formed the merged 1240K dataset and the merged HO dataset (**Table S1**). In the merged higher-density Illumina dataset using for haplotype-based analysis, we merged genome-wide data of Miao with our recent publication data from Han, Mongolian, Manchu, Gejia, Dongjia, Xijia and others (Chen et al., 2021a; He et al., 2021; Liu, Y. et al., 2021b; Yao et al., 2021).

### Frequency-based population genetic analysis

#### Principal component analysis

We performed principal component analysis (PCA) in three population sets focused on a different scale of genetic diversity. Smartpca package in EIGENSOFT software (Patterson et al., 2006) was used to conduct PCA with ancient sample projected and no outlier removal (numoutlieriter: 0 and lsqproject: YES). East-Asian-scale PCA included 393 TK people from 6 Chinese populations and 21 Southeast populations; 144 HM individuals from 7 Chinese populations and 6 Southeast populations; 968 Sinitic people from 16 Chinese populations, 356 TB speakers from 18 northern and 17 southern populations, 248 AA people from 20 populations, 115 Austronesian (AN) people from 13 populations, 304 Trans-Eurasian people from 27 populations from North China and Siberia and 231 ancient individuals from 62 groups. Chinese-scale PCA was conducted based on the genetic variations of Sinitic, northern TB and TK people in China, ancient populations from Guangxi, and all 16 HM-speaking populations. Twenty-three ancient samples from 9 Guangxi groups were projected (Wang, T. et al., 2021). The third HM-scale PCA included 15 modern populations (Vietnam Hmong populations show as outliers) and two Guangxi ancient populations.

#### ADMIXTURE

We performed model-based ADMIXTURE analysis using the maximum likelihood clustering in the ADMIXTURE (Version 1.3.0) software (Alexander et al., 2009) to estimate individual ancestry composition. Included populations in the East Asian-scale PCA analysis and Chinese-scale PCA analysis were used in the two different AMIDXUTRE analyses with the respective predefined ancestral sources ranging from 2 to 16 and ranging from 2 to 10. We used PLINK (version v1.90) to prune the raw SNP data into unlinked data via pruning for high linkage disequilibrium (−indep-pairwise 200 25 0.4). We estimated the cross-validation error using the results of 100 times ADMIXTURE runs with different seeds. And the best-fitted admixture model was regarded being possessed the lowest error.

#### Phylogeny modeling with TreeMix

We used PLINK (version v1.90) to calculate the pairwise Fst genetic distance between studied Sichuan Miao (SCM) and other modern and ancient references and also estimated the allele frequency distribution of included populations in the TreeMix analyses. Both modern and ancient populations were used to construct the maximum-likelihood-based phylogenetic relationship with population splits and migration events using TreeMix v.1.13 (Pickrell and Pritchard, 2012).

#### Outgroup-*f_3_*-statistics and admixture-*f_3_*-statistics

We assessed the potentially existed admixture signatures in SCM via the admixture-*f_3_*-statistics in the form *f_3_*(Source1, source2; Miao_Baila/Jiancao/Hele), which was calculated using qp3Pop (version 435) package in the AdmixTools software (Patterson et al., 2012). The target populations with the observed negative *f_3_* values and Z-scores less than −3 were regarded as mixed populations with two surrogates of ancestral populations related to source1 and source2. Followingly, similar to the quantitation of the genetic similarities and differences as pairwise Fst, we assessed the genetic affinity between studied populations and other reference populations via the outgroup - *f_3_*-statistics in the form of *f_3_*(Reference source, studied Miao; Mbuti).

#### Pairwise qpWave tests

We calculated p-values of the rank tests of all possible population pairs among HM-speaking populations and other geographically close modern and ancient reference populations using qpWave in the AdmixTools package (Patterson et al., 2012) to test their genetic evolutionary relationships and genetic homogeneity. Here, we used a set of distant outgroup sets, which included Mbuti, Ust_Ishim, Kostenki14, Papuan, Australian, Mixe, MA1, Jehai and Tianyuan. The obtained pairwise matrix of the p values was visualized and presented in a heatmap using pheatmap package.

#### Admixture modeling using qpAdm

We further assessed the relative ancestral source and corresponding admixture proportion of Chinese HM-speaking and surrounding Han Chinese populations using a two-way-based admixture model in the qpAdm (version 634) in the AdmixTools package (Patterson et al., 2012). One of the studied populations combined with two predefined ancestral modern and ancient sources were used as the left populations and the aforementioned pairwise-based outgroups were used as the left populations along with two additional parameters (allsnps: YES; details: YES).

#### Demographic modeling with qpGraph

We used the R package of ADMIXTOOLS2 (Patterson et al., 2012) to explore the best-fitted phylogenetic topology with admixture events and mixing proportions with the Mbuti, Onge, Loschbour, Tianyuan, Baojianshan, Qihe, GaoHuaHua, Longshan as the basic representative genetic linages for molding the formation of modern SCM. A “rotating” scheme of adding other modern and ancient populations was used to explore other genetic ancestries that would improve the qpGraph-based admixture models. One model with the predefined admixture events ranging from 0 to 5 was ran 50 times and we then choose the best models based on the Z-scores and best-fitted scores. We also replaced the Longshan people with the upper Yellow River Lajia people as the northern ancestral lineage and ran all aforementioned admixture models.

#### ALDER

We estimated the decay of linkage disequilibrium in SCM using all possible population pairs of modern East Asians as surrogate populations in ALDER 1.0 (Loh et al., 2013). Two additional parameters were used here: jackknife: YES and mindis: 0.005.

### Haplotype-based population genetic analysis

#### Segmented haplotype estimation

We used Shapeit software (Segmented HAPlotype Estimation & Imputation Tool) to phase our dense SNP data with the default parameters (--burn 10 --prune 10 --main 30) (Browning and Browning, 2011). Pairwise sharing IBD segments were calculated using refined-ibd software (16May19.ad5.jar) with the length parameter as 0.1 (Browning and Browning, 2013).

#### Chromosome painting

We ran ChromoPainterv2 software (Lawson et al., 2012) to paint the target SCM and sampled surrogate northern and southern East Asians using all phased populations as the surrogate populations, which was regarded as the full analysis. We also removed the SCM and their most close genetic relatives (Gejia, Dongjia and Xijia) in the set of surrogates and painted all target and surrogate populations once again, which was regarded as the regional analysis. We then combined all chunk length output files of 22 chromosomes as the final dataset of sharing chunk length.

#### FineSTRUCTURE analysis

we identified the fine-scale population substructure using fineSTRUCTURE (version 4.0) (Lawson et al., 2012). Perl scripts of convertrecfile.pl and impute2chromopainter.pl were used to prepare the input phase data and recombination data. FineSTRUCTURE, ChromoCombine and ChromoPainter were combined in the four successive steps of analyses with the parameters (-s3iters 100000 -s4iters 50000 -s1minsnps 1000 -s1indfrac 0.1). The estimated coancestry was used to run PCA analysis and phylogenetic relationships at the individual-level and population-level.

#### GLOBETROTTER-based admixture estimation

We ran the R program of GLOBETROTTER (Hellenthal et al., 2014) to further identify, date and describe the admixture events of the target SCM. Both painting samples and copy vectors estimated in the ChromoPainterv2 were used as the basal inputs in the GLOBETROTTER-based estimation. We first ran it to infer admixture proportions, dates and sources with two specifically predefined parameters (prop.ind: 1; bootstrap.num:20) and we then reran it with 100 bootstrap samples to estimate the confidence interval of the admixture dates.

#### Natural selection indexes of XPEHH and iHS estimation

We calculated the integrated haplotype score (iHS) and cross-population extended haplotype homogeneity (XPEHH) using the R package of ReHH (Gautier et al., 2017). Here both northern Han Chinese from Shaanxi and Gansu provinces, and southern Han Chinese from Sichuan, Chongqing and Fujian provinces were used as the reference in the XPEHH estimation.

#### Gene enrichment analysis

The online tool of Metascape (Zhou et al., 2019) was used to annotate the potentially existed natural selection signatures in the iHS and XPEHH.

## Results

### Newly-identified HM genetic cline in the context of East Asian populations

We genotyped 52 genome-wide SNP data in three SCM populations (Baila, Jiancao and Hele) and found that five samples possessed close sibship with other samples. After removing relatives, we merged our data with the Human Origin dataset in AADR (merged HO dataset) to explore the genetic diversity of SCM and their genetic relationship with modern and ancient Eurasian populations. East-Asian-scale PCA results showed three genetic clines (**Figure 1A**), which included the northern East Asian cluster (Altaic and northern ST speakers), and the southern East Asian and Southeast Asian cluster (AA, AN, TK and southern TB), and the newly identified HM genetic cline. Interestingly, our newly-studied three SCM populations separated from other Chinese populations and clustered closely with geographically distant Hmong people from North Vietnam (Hmong) and Thailand (Hmong-Daw and Hmong-Njua), suggesting their strong genetic affinity and potentially existing common origin history. Dao and IuMien clustered closely with TK people, and Miao and She people from Chongqing and other southern China were overlapped with geographically close Han people, which suggested the massive population interaction between HM people and their neighbors. Other HM people, including Geijia, Dongjia and Xijia in Guizhou, PaThen in Vietnam were localized between three genetically different HM genetic lineages.

**Figure 1.**
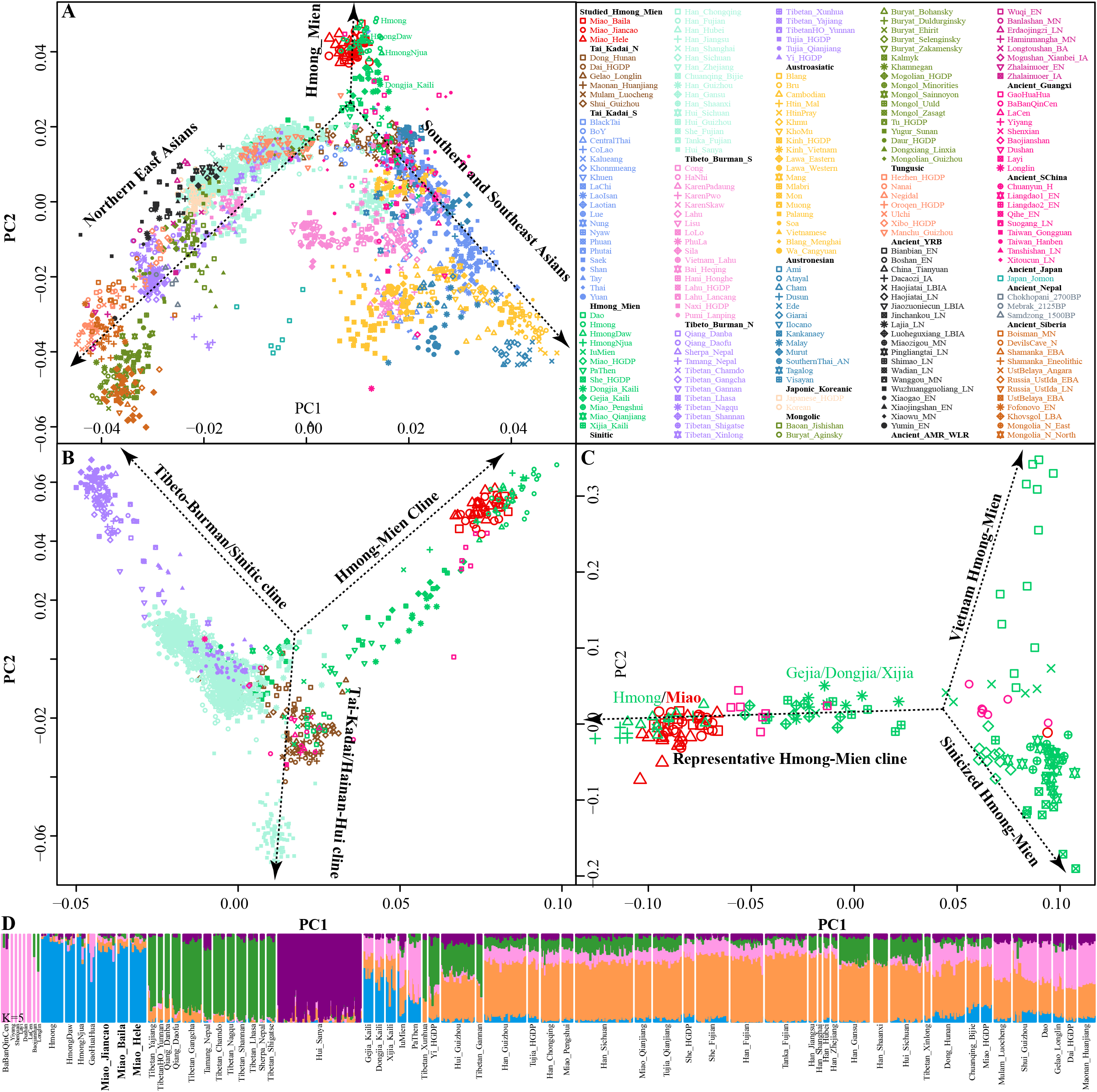
Genetic affinity of HM people in the context of modern and ancient eastern Eurasians. (**A~C**), Principal component analyses focused on the genetic diversity from East Asian, South Chinese populations and HM-speaking populations. Included East Asian ancient populations were projected onto the modern genetic background. Populations were color-coded based on geographical and linguistic categories. (**D**), Model-based ADMIXTURE results with five predefined ancestral sources showed the ancestral cluttering pattern and their individual ancestral proportion.

Focused on the genetic diversity of ST and TK people in China and all studied and reference HM populations, we used a panel of 65 populations and identified three primary directions in the first two dimensions represented by ST, HM and Hainan Hui people [(top right, top left, and bottom, respectively), **Figure 1B**]. We found that ~500-year-old prehistoric Guangxi GaoHuaHua was localized closely with SCM, but ~1500-year-old BaBanQinCen overlapped with Chinese TK people and HM Dao. Additionally, we explored the finer-scale population relationship within geographically different Miao populations and found that Vietnam Hmong separated from other populations along PC2. After removing this outlier of Hmong, PCA patterns also showed three different genetic clades among the remaining sixteen HM populations, which represented by representative HM cline, Sinicized HM and Vietnam HM [(right, top left, and bottom left, respectively), **Figure 1C**]. These identified population stratifications among HM-speaking populations were confirmed via pairwise Fst genetic distances among 29 Chinese populations based on the Illumina-based dataset (**Table S2**) and among 65 populations based on the merged HO dataset (**Table S3**). Genetic differences estimated via Fst values showed that SCM had a close genetic relationship with Guizhou HM people (Gejia, Dongjia and Xijia), followed by geographically different ST groups, northern Mongolic Mongolian and southern AA populations (Blang and Wa). Results from the lower-density HO dataset not only confirmed the general patterns of genetic affinity between SCM and East Asians reported in Illumina-dataset but also directly identified that SCM possessed the genetic affinity with Hmong people from Vietnam and Thailand among modern reference populations, with GaoHuaHua (Miao_Baila: 0.1398; Miao-Jiancao: 0.1394; Miao_Hele: 0.1419) among ancient Guangxi ancient references.

### Ancestral composition of HM-speaking populations

Consistent with the identified unique genetic cluster of SCM people, we expectedly observed one dominant unique ancestry component in HM-speaking populations (blue ancestry in **Figure 1D**). HM-specific ancestry maximized in Vietnam and Thailand Hmong people, as well as existed in SCM and GaoHuaHua with a higher proportion. Different from the gene pool of HM people in Southeast Asia, SCM and ~500-year-old GaoHuaHua people harbored more ancestry related to 1500-year-old historic Guangxi people (pink ancestry). Furthermore, SCM harbored more genetic influence from Sinitic-related populations (origin and purple ancestries) relative to the GaoHuaHua people. A similar pattern was observed in Guizhou populations but with different ancestry proportions, in which Guizhou HM people harbored higher pink and orange ancestries and smaller blue ancestry. This observed pattern of ancestry composition suggested that Guizhou and Sichuan HM-speaking populations absorbed additional gene flow from northern East Asians when they experienced extensive population movement and interaction. Indeed, other Miao people from Chongqing and She and Miao in the HGDP project possessed similar ancestry composition with neighboring Hans, which supported the stronger extent of admixture between Pro-HM and incoming southward Han’s ancestor. The admixture signatures in the *f_3_*(East Asians, Miao_Baila; Miao_Jiancao) confirmed that Jiancao Miao was an admixed population and harbored additional genetic materials from northern East Asians (negative Z-scores in LateXiongnu (−3.798), LateXiongnu_han (−3.506), Han_Shanxi (−3.076) et.al.) and southern East Asians (−3.443 in Li_Hainan) (**Table S4**). However, no statistically significant negative *f_3_*-values have been identified in the targets of the other two SCM groups. Evidence from the ancient genomes has suggested that prehistoric Guangxi GaoHuaHua people were the temporally direct ancestor of modern Guangxi Miao people (Wang, T. et al., 2021). However, only marginal negative *f_3_*-values were observed in Jiancao Miao, as *f_3_*(GaoHuaHua, Pumi_Lanping; Miao_Jiancao)=-1.228*SE, although we observed a close cluster relationship in the PCA and ADMIXTURE.

To further characterized the admixture landscape of SCM and other East Asian representative populations based on the sharing haplotypes, we used SCM as the surrogate of the ancestral source and painted all other sampled East Asian populations using ChromoPainter. We found Guizhou HM populations (Gejia, Dongjia and Xijia) copied the longest DNA chunk from SCM with the total copied chunk length over 1287.74 centimorgans (**Figure 2A**). SCM also contributed much genetic material to geographically close Miao, Han and Chuanqing groups (over 237.31 centimorgans) and donated relatively less ancestry to northern Altaic- and southern AA and TB-speaking populations, including the Wa, Pumi, Lahu and Bai in geographically close Yunnan Province (**Figure 2B**). Followingly, we explored the extent to which other putative East Asian surrogates contributed to the formation of the SCM people. We used other non-HM people as the ancestral surrogate to paint the SCM people and we found southern Han Chinese donated much ancestry to targeted Miao (**Figure 2C**), even higher than southern Miao and She and other southern East Asian indigenous populations (**Figure 2D**), which provided supporting evidence for genetic interactions between HM and southern Sinitic people. Collectively, the SCM people served as one unique ancestral source that contributed much genetic ancestry to modern East Asians.

**Figure 2.**
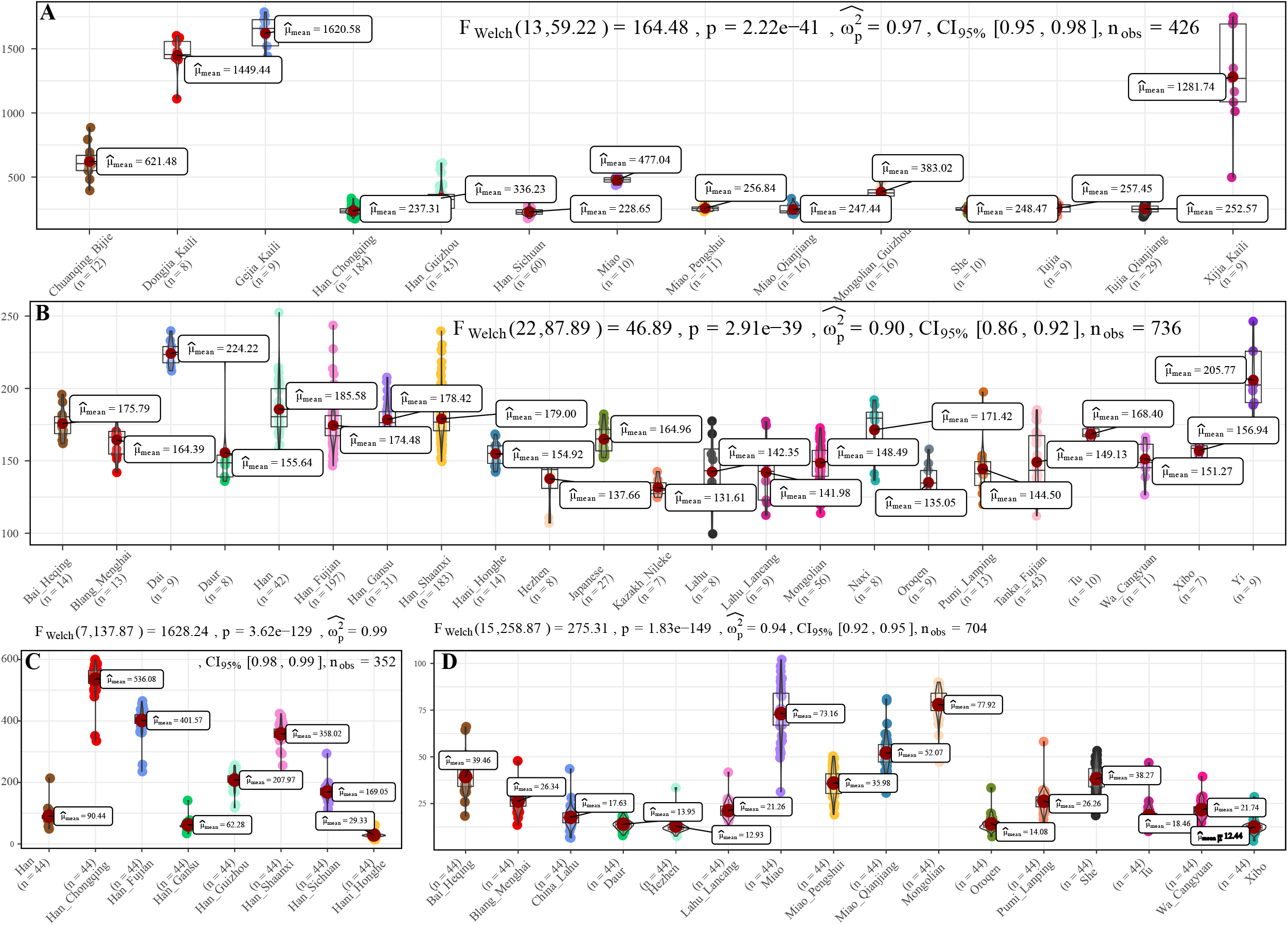
The chromosome painting between Hmong-Mien people and other East Asian reference populations. (**A~B**), the amount of total length of DNA fragments of modern East Asians copied from donor chromosomes from Sichuan Miao. (**C~D**), The average DNA chunk of Sichuan Miao copied from other East Asians. Statistical indexes showed the results of inter-population comparisons

Although the genetic affinity between SCM and Sinitic Han Chinese were identified, finer-scale population structure inferred from the fineSTRUCTURE showed that SCM possessed a similar pattern sharing ancestry with Guizhou HM people and formed one specific HM branch (**Figure 3**). The inferred PCA patterns based on the sharing haplotypes showed that SCM separated from other Han Chinese and Yunnan AA and TB people and had a close relationship with Guizhou HM people (**Figure 3A~C**). Clustering patterns based on the sharing DNA fragments among population-level and individual-level (**Figure 3D~E**) further confirmed the genetic differentiation between HM people and Sinitic people, which is consistent with the genetic affinity observed in the shared IBD matrix. Additionally, we used the GLOBETROTTER to identify, date and describe the admixture status of SCM. We first conducted the regional analysis, in which meta-SCM was used as the targeted populations and other East Asians except to Guizhou HM people used as the surrogates. The best-guess conclusion was an unclear signal, which provided evidence for their unique population history of SCM. Thus, we secondly performed full analysis to characterized three SCM people conditional on all other sampled East Asian populations as ancestral proximity. We identified recent admixture events in all three geographically different targets. One-date admixture model for Baila Miao suggested that it was formed via recent admixture events in seven generations ago with one source related to Jiancao Miao (0.86) and the other source related to Sichuan Han (0.14). A similar admixture model was identified in Hele Miao people, in which the identified one-date model showed that a recent admixture event occurred five generations ago with major ancestry sources related to Jiancao Miao (0.84) and the minor source related to Guizhou Han (0.16). We found two-date-two-way admixture model best fitted the genetic admixture history of Jiancao Miao. The ancient admixture events occurred 86 generations ago with the Guizhou Gejia as the minor source proximity (0.48) and Baila Miao as the major source proximity (0.52). A recent admixture occurred five generations ago with Baila Miao as the major donor (0.83) and Guizhou Han as the minor donor (0.17). We further estimated the admixture times used ALDER using three SCMs as the targets and all other modern East Asians as the ancestral sources to test the decay of linkage disequilibrium (**Table S5**). When we used Guizhou HM people as one of the sources, both population compositions from northern and southern East Asians can produce statistically significant admixture signatures with the admixture times ranged from 22.35+/-6.92 (Maonan) to 160.58±70.32 (Xijia), which also provided supporting clues for the complex ancient admixture events for different ancestral sources.

**Figure 3.**
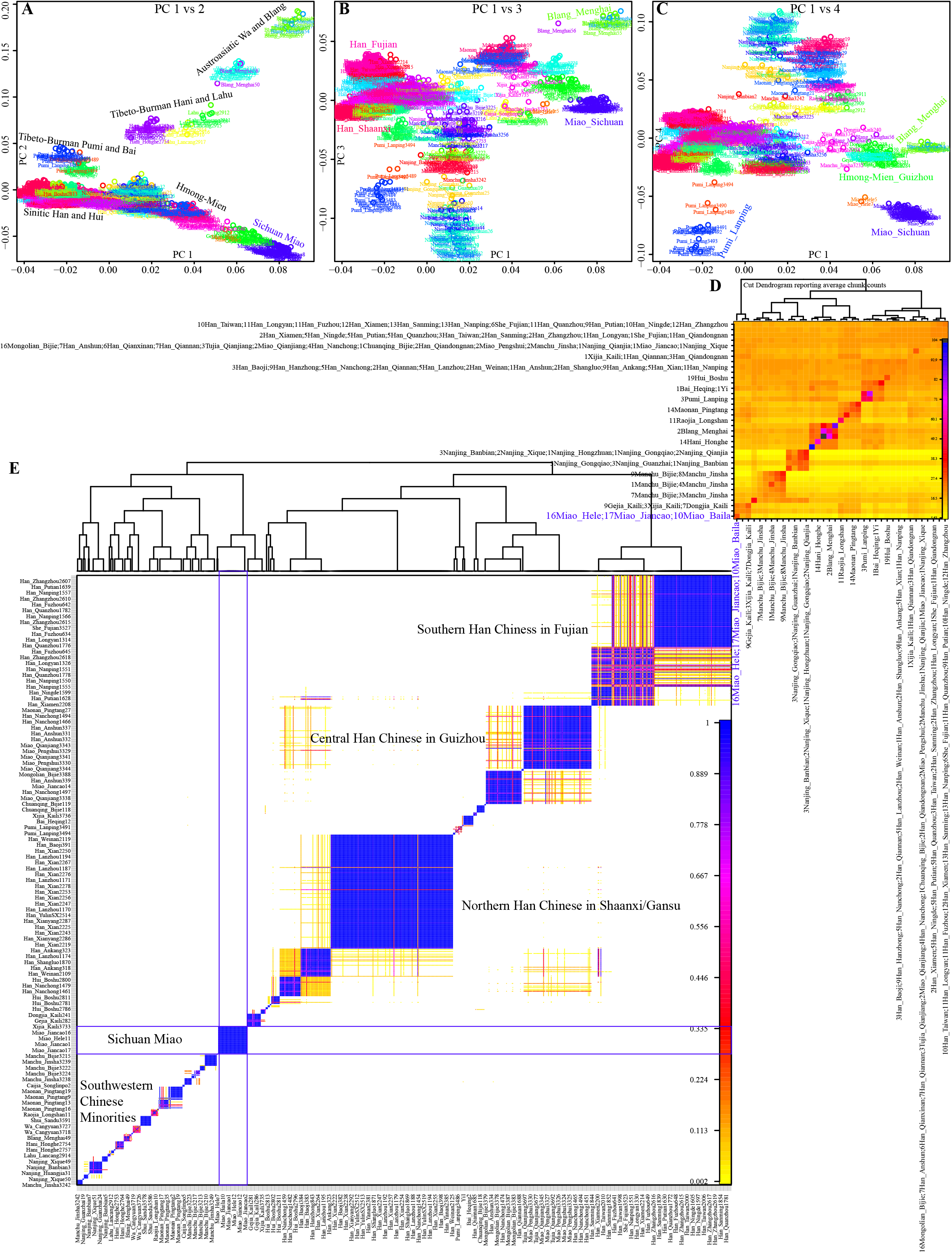
Fine-scale population genetic structure based on the shared haplotype data. (**A~C**), PCA results based on the co-ancestry matrix showed a genetic relationship among modern East Asians. The color showed the re-classification of the homogeneous population label. (**D~E**), clustering patterns of individual-level and population-level East Asians based on the pairwise coincidence matrixes.

### Genetic admixture and continuity of HM-specific ancestry at the crossroads of East and Southeast Asia in the past 1500 years

To further explore the geographic distribution of our identified HM-dominant ancestry and further constrain the formed time range, we conducted a series of formal tests to validate our predefined phylogenetic topologies. Shared genetic drift inferred from outgroup-*f_3_*-statistics in the form *f_3_*(SCMs, modern East Asians; Mbuti) suggested that SCM shared a closest genetic relationship with Guizhou HM people, followed by TK people in South China and geographically close Han based on the merged 1240K dataset (**Table S6**). The genetic affinity between SCM and Hmong people in Vietnam and Thailand was directly evidenced via the observed largest outgroup-*f_3_*-values in the merged HO dataset, suggesting HM-specific ancestry widely distributed in Sichuan, Guizhou, Guangxi, Vietnam and Thailand. Focused on the ancient reference populations, we found that historic Guangxi GaoHuaHua people were on the top list for the shared genetic drift (0.3324 for Baila Miao, 0.3317 for Hele Miao and 0.3304 for Jiancao Miao). 1500-year-old Guangxi BaBanQinCen and Iron Age Taiwan Hanben also possessed strong genetic affinity with SCM, suggesting their common origin history and possibly originated from South China. These patterns of genetic affinity among spatiotemporally different southern East Asians were consistent with the shared characteristics attested by cultural, linguistic and archeological documents.

To further explore the genetic relationship between ancient Guangxi populations and modern ethnolinguistic populations, we conducted pairwise qpWave analysis among 16 HM populations, five Guangxi ancient groups (GaoHuaHua, BaBanQinCen, Baojianshan, Dushan and Longlin), seven TK-, 16 Sinitic- and 18 TB-speaking populations (**Figure 4**). We found genetic homogeneity existed within populations from geographically and linguistically close populations, especially in TB, Sinitic and HM. Here, we only observed strong genetic affinity within geographically diverse HM people and found genetic heterogeneity between historic Guangxi populations and modern HM people. Considering different admixture models identified among three SCM populations, we performed symmetrical *f_4_*-statistics in the form *f_4_* (SCM1, SCM2; reference populations, Mbuti) (**Table S7**). We also identified differentiated evolutionary history among them; Jiancao Miao shared more alleles with Guizhou HM people compared with Miao people from Baila and Hele, and Jiancao Miao also shared more northern East Asian ancestry related to other two Miao populations. Results from another version of symmetrical *f_4_*-statistics in the form *f_4_*(reference1, reference2; SCM, Mbuti) first confirmed the strong genetic affinity between SCM people and other HM people, as most negative *f_4_*-values identified in *f_4_*(reference1, HM; SCM, Mbuti) (**Table S8**). All 126 tested *f_4_*(Reference, GaoHuaHua; SCM, Mbuti) values were negative and 123 out of 126 were statistically significant, which suggested the SCM shared more ancestry and a closer genetic relationship with GaoHuaHua relative to other modern and ancient East Asians. We al so tested *f_4_*(Reference, SCM; GaoHuaHua, Mbuti) (**Table S9**) and found GaoHuaHua shared more alleles with SCM compared to all reference populations. These observed results were consistent with the hypothesis of SCM people are the direct descendants of historic Guangxi GaoHuaHua. We also tested *f_4_*(GaoHuaHua, SCM; reference, Mbuti) and found additional gene flow from ancestral sources related to late Neolithic populations from the YRB, as observed negative *f_4_*-values in *f_4_*(GaoHuaHua, Miao_Hele; Han_Gansu, Mbuti)=-3.78*SE or *f_4_*(GaoHuaHua, Miao_Baila; China_Upper_YR_LN, Mbuti)=-3.252*SE. Indeed, we previously observed admixture signatures in Jiancao Miao in admixture-*f_3_*(GaoHuaHua, northern East Asians; Jiancao Miao), which suggested SCM shared major ancestry from GaoHuaHua and also experienced additional genetic admixture from northern East Asians.

**Figure 4.**
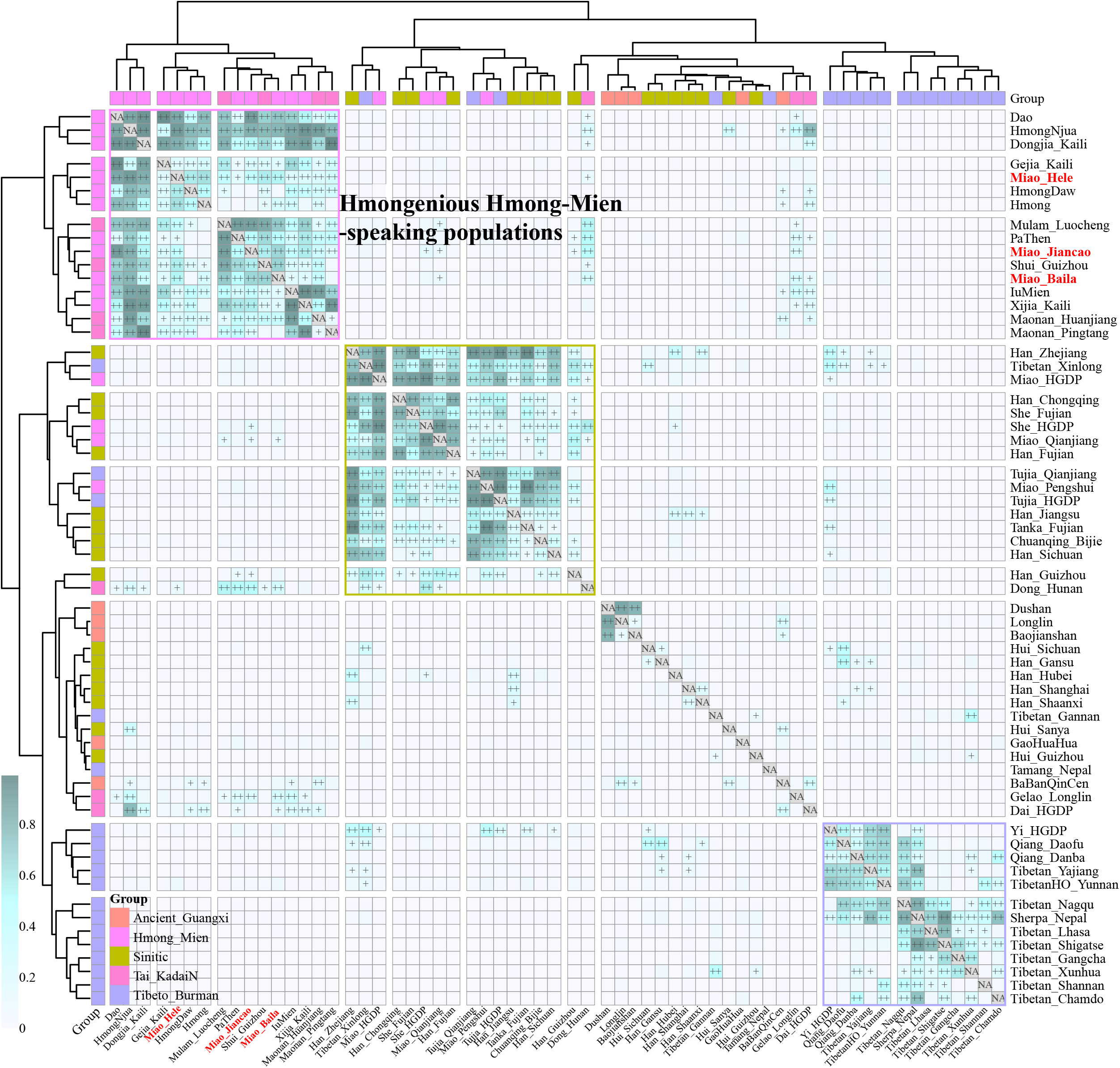
Pairwise qpWave analysis showed the genetic heterogeneity and homogeneity among East Asians. P-values of rank1 tests larger than 0.05 showed the genetic homogeneity among two reference populations, which were marked as “++”, and p values of rank1 tests larger than 0.01 were marked as “+.”

Focused on the deeper temporal population dynamics, we followingly tested the genetic relationship between SCM and ~1500-year-old BaBanQinCen used the same strategies (**Table S9**). Positive results in *f_4_*(Dongjia/Maonan/China_SEastAsia_Coastal_LN/ Guangxi_1500BP, SCM; BaBanQinCen, Mbuti) showed that BaBanQinCen shared more derived alleles with late Neolithic and Iron Age Fujian populations and other spatiotemporally close Guangxi historic populations. Statistically significant values in *f_4_*(BaBanQinCen, SCM; reference, Mbuti) further confirmed that BaBanQinCen does not form a clade with SCM and shared more alleles with pre-Neolithic Amur River people (AR14K), Neolithic-to-Iron Age Fujian populations and indigenous Guangxi prehistoric populations (Baojianshan and Dushan) compared with SCM, which was further supported via the *f_4_*-statistics focused on other ~1500-year-old Guangxi populations (Guangxi_1500BP) and Taiwan Hanben. But SCM shared more genetic influence from northern East Asians compared with ~1500-year-old Guangxi people. Compared with other Guangxi prehistoric populations (*f_4_*(Longlin, Baojianshan and Dushan, reference; SCM, Mbuti)), SCM shared more ancestry with ancient northern East Asians, southern Fujian and modern East Asian ancestry. Compared with SCM, prehistoric Guangxi populations shared more Neolithic to Iron Age Fujian and Guangxi ancestries. We also tested the genetic relationship between SCM and YRB farmers using asymmetric-*f_4_*-statistics and found YRB millet farmers shared more alleles with SCM people compared to early Asians and southern Fujian and Fujian ancient populations. As expected, SCM harbored more HM-related alleles or ancient Fujian and Guangxi ancestries compared with millet farmers. Generally, formal test results demonstrated that SCM possessed the strongest genetic affinity with ~500 - year-old Guangxi GaoHuaHua people and additionally obtained genetic influx from northern East Asians recently.

### Admixture evolutionary models

A close genetic relationship between Guangxi historic populations and SCM has been evidenced in our descriptive analyses and quantitative *f*-statistics. We further conducted two-way qpAdm models with two Guangxi ancient populations as the southern surrogates and four northern ancient populations from YRB and Amur River as the northern ancestral sources to estimate the ancestral composition of SCM and their ethnically and geographically close populations (**Figure 5A**). When we used BaBanQinCen as the source, we tested the two-way admixture models: proportion of ancestry contribution of historic Guangxi population ranged from 0.811±0.107 in Kali Dongjia to 0.404±0.107 in Shaanxi Hans in the AR14K-BaBanQinCen model and spanned from 0.738±0.145 to 0.127±0.088 in Shaanxi Hans in China_YR_LBIA-BaBanQinCen model. SCM derived 0.780~0.806 ancestry from historic Guangxi ancestry in the former model and 0.653~0.666 ancestry from it in the latter model (**Figure 5A**). We also confirmed that the unique gene pool of SCM derived from major ancestry from Guangxi and minor ancestry from North East Asians via the additional two qpAdm admixture models with early Neolithic Amur River Hunter-Gatherer, middle Neolithic-to-Iron Age YRB farmers as the northern sources.

**Figure 5.**
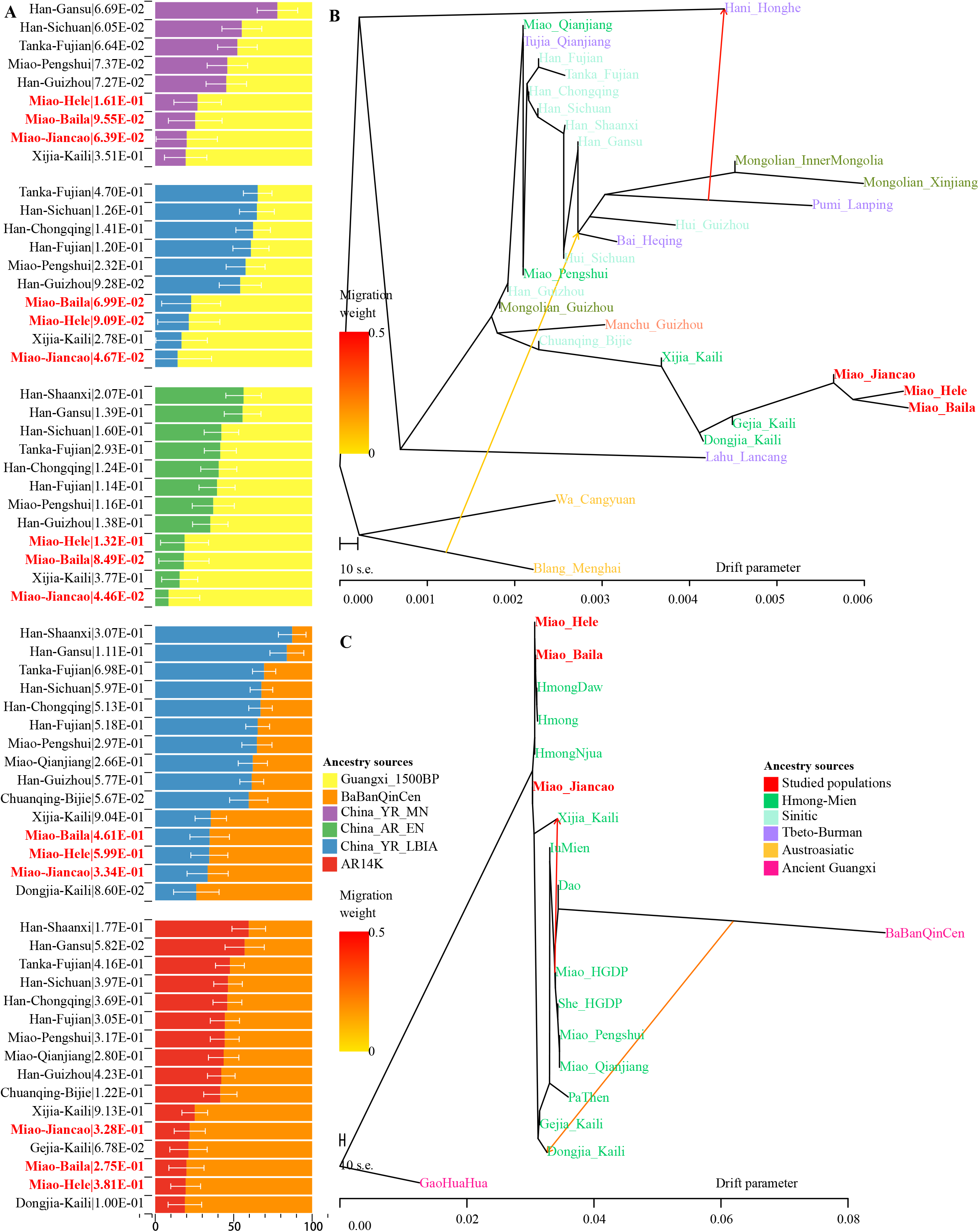
Results of qpAdm models and TreeMix-based phylogenies. (**A**), Two-way admixture models showed ancestry comparison in different ancestral source pairs. (**B~C**), TreeMix-based phylogenetic tree with two migration events showed the geneti c relationship between East Asians.

Until now, to explore the population genetic diversity of Chinese populations and provided some pilot works supporting the initiation of the Chinese population genome diversity project (CPGDP) based on the deep whole-genome sequencing on anthropologically-informed sampling populations, we have genotyped the array-based genome-wide SNP data in 29 ethnolinguistically different populations. We reconstructed phylogenetic relationships between three studied SCM populations and 26 other Chinese populations from ST, Altaic, AA and HM (**Figure 5B**). We identified that branch clusters were consistent with the linguistic categories and geographical division. Tibetan Lahu and Hani clustered closely with AA Blang and Wa, and other populations were clustered as the northern and southern East Asian branches. The southern branches consisted of our newly-studied Miao and Guizhou HM people and Guizhou Chuanqing and Manchu. The northern branch comprised of Mongolic, TB and Sinitic people. We found that two Chongqing Miao populations clustered closely with the northern branch, suggesting much genetic material mixed from surrounding Han Chinese populations. We also identified regional population gene flow events from ethnically different populations, such as gene flow events from Pumi to Hani and from Blang to common ancestral lineage of Bai, Pumi and Mongolian. To directly reconstruct the phylogenies between the HM population and historic Guangxi populations, we merged 16 HM-speaking populations with GaoHuaHua and BaBanQinCen and found two separated branches respectively clustered closely with GaoHuaHua and BaBanQinCen (**Figure 5C**). Close phylogenetic relationship among SCM, Guangxi GaoHuaHua, Guizhou Gejia, Dongjia and Xijia, and Vietnam and Thailand Hmong further supported the common origin of geographically different HM people.

We finally reconstructed the deep population admixture history of HM-speaking populations using the qpGraph model with population splits and admixture events. We used the ancestral lineage of Mbuti in Africa, Loschbour in western Eurasia, Onge in South Asia, Tianyuan in East Asia as the basal deep early continental lineages. We used Baojianshan in the early Neolithic period and GaoHuaHua in the historic time from Guangxi, Qihe in the early Neolithic in Fujian as southern East Asian lineages and used Neolithic YRB millet farmers as the northern East Asian lineages. In our first best-fitted model (**Figure 6A**), we added additional late Fujian Xitoucun and Tanshishan from the late Neolithic period, we found GaoHuaHua could be fitted as major ancestry related to upper Yellow River Qijia people (0.52) and minor ancestry related to late Neolithic Fujian people (0.48). However, SCM derived much more ancestry from northern East Asians (0.82) in this model, suggesting additional northern East Asian gene flow influenced the genetic formation of modern HM-speaking populations. In the second bested fitted model (**Figure 6C**), we added Hunter-Gatherer lineage from the Mongolian Plateau (Bosiman) and found Xuyong Miao could be fitted as 0.86 ancestry from GaoHuaHua the remaining ancestry from Qijia people (0.14). The third best-fitted model (**Figure 6D**) with adding Australian lineage also replicated the shared major ancestry between GaoHuaHua and Xuyong Miao. In the final version of the qpGraph model (**Figure 6D**), we added the American indigenous lineages, in which Miao was fitted as 0.37 ancestry from western Eurasian and 0.63 ancestry from East Asians. Xuyong Miao was modeled as similar ancestry composition as the third model. Here, we should be cautious that the differences in the topologies of the early deep lineages when different populations were added to our basal models. The detailed true phylogenetic relationship should be further explored and reconstructed via denser spatiotemporally different early Asian population sequencing data. But the consistent pattern of Miao’s genetic profiles of major ancestry from GaoHuaHua and minor ancestry from northern East Asia were obtained from four different admixture models, suggesting it is valuable to illuminate the simple model of the formation of modern SCM.

**Figure 6.**
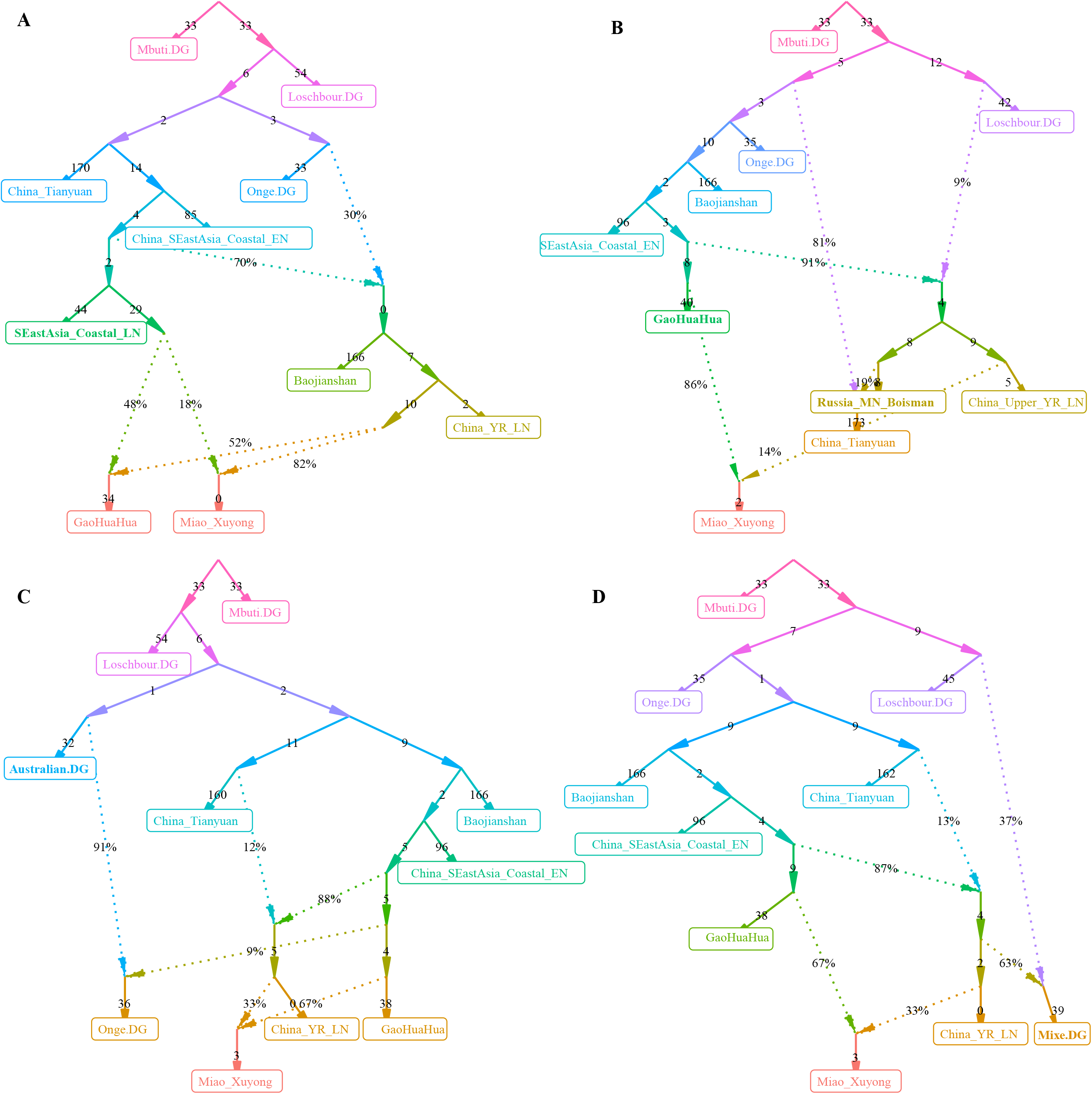
Deep population history reconstruction based on the best-fitted qpGraph models. Different frameworks in the qpGraph-based models adding the late Neolithic Fujian population (SEastAsia_Coastal_LN, **A**), Mongolian Plateau Hunter-Gatherer (Boisman, **B**), Australian (**C**) and Mixe (**D**).

### Uniparental founding lineages

We obtained high-resolution uniparental maternal and paternal lineages in SCM (**Table S10**). We identified four dominant maternal founding lineages in SCM [(B5a1c1 (0.3462), F1g1 (0.1346), B4a (0.0769) and F1a (0.0769)]. We also identified two paternal founding lineages [(O2a2a1a2a1a2 (0.3913) O2a1c1a1a1a1a1a1b (0.1739)] in SCM, which is consistent with the hypothesis of the primary ancestry of Miao originated from southern Chinese indigenes. In detail, we observed 10 terminal paternal lineages among 23 males and 17 terminal maternal lineages in 52 females. Compared with geographically close Chongqing Han populations, we found a significant difference in the frequency of major lineages between Chongqing Han and Sichuan Miao (**Figure 7**).

**Figure 7.**
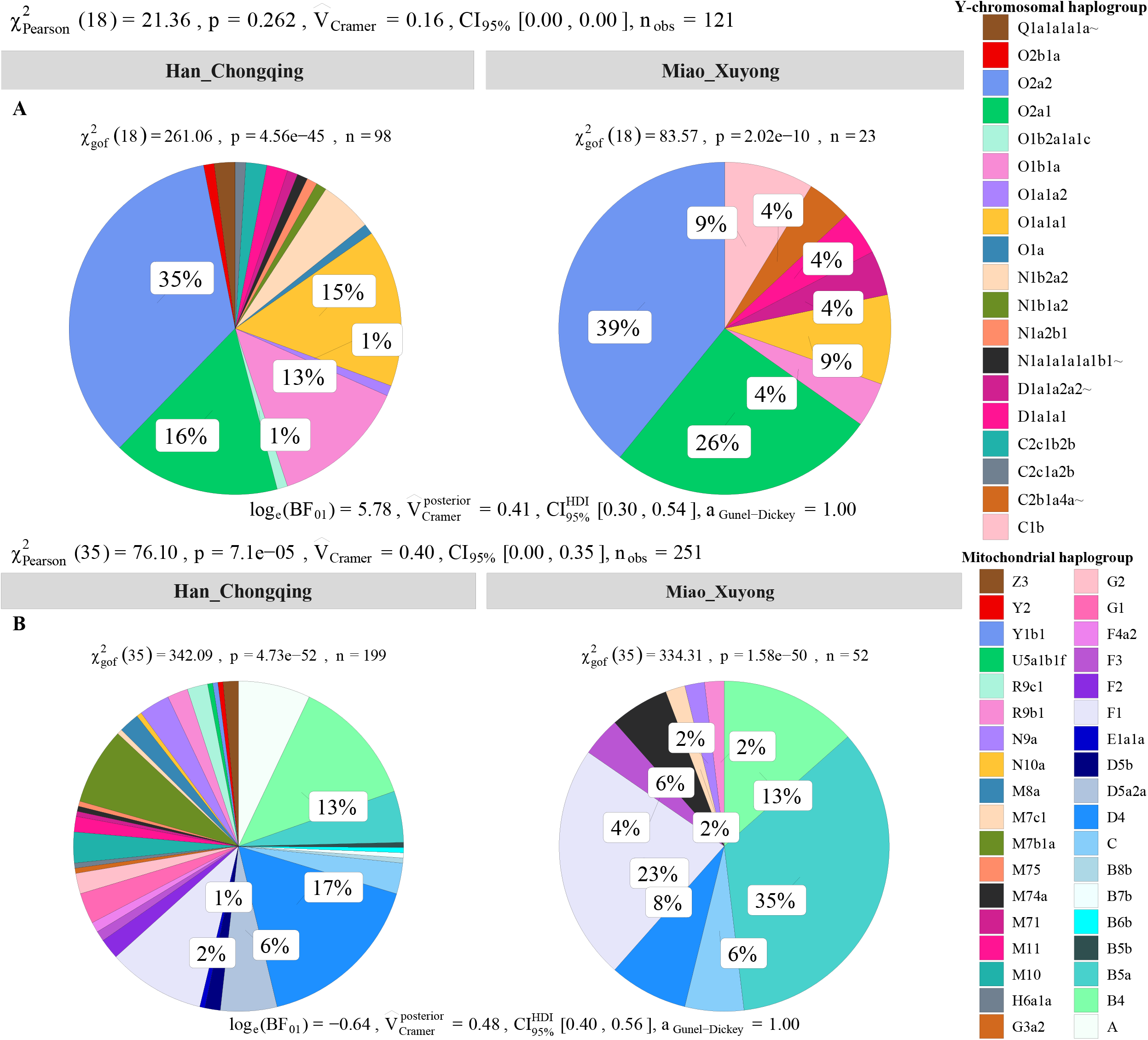
Allele frequency spectrum of observed maternal and paternal haplogroups of Chongqing Miao and Chongqing Han. Population comparison between Han and Miao based on the frequency distribution of the observed paternal lineages (**A**) and maternal lineages (**B**) via the Pearson and Cramer tests.

### Natural selection signatures and their biological adaptation

Genetic studies have identified many biologically adaptive genes or pathways in ethnoling uistically diverse populations. Evolutionary adaptative mutations could be accumulated and generated longer extended haplotype homozygosity with their increase of allele frequency of the initial mutations. We scanned for candidates of the positive selections using iHS and XPEHH in SCM. We first calculated XPEHH values for Miao using northern Han as reference population and identified obvious candidates in Chromosomes 1-3, 9, 20 and 22 (**Figure 8A**). Chromosome 1 showed selection signals in the vicinity of *Neuroblastoma breakpoint family member 9/10* (NBPF 9/10) locus, reflecting well-known signals associated with susceptibility of the neuroblastoma. We further identified a strong selection signal implicating *Polypeptide N-acetylgalactosaminyltransferase 13* (GALNT13) and *potassium voltage-gated channel subfamily J member 3* (KCNJ3) located in chromosome 3. The former one is expressed in all neuroblastoma cells and encodes a glycosyltransferase enzyme responsible for the synthesis of O-glycan. The latter one encodes G proteins in the potassium channel and is associated with susceptibility candidates for schizophrenia (Yamada et al., 2012). We also identified four top candidate genes in chromosome 3, including the *abhydrolase domain containing 10* (ABHD10), *RNA binding motif single stranded interacting protein 3* (RBMS3), *RBMS3 antisense RNA 3* (RBMS3-AS3) and *transgelin 3* (TAGLN3). ABHD10 is one of the important members of the AB hydrolase superfamily and is associated with enzymes for deglucuronidation of mycophenolic acid acyl-glucuronide (Iwamura et al., 2012). RBMS3 encodes protein binding Prx1 mRNA in a sequence-specific manner via binding poly(A) and poly(U) oligoribonucleotides and controls Prx1 expression and indirectly collagen synthesis (Fritz and Stefanovic, 2007). It also served as the tumor suppressor gene associated with lung squamous cell carcinoma and esophageal squamous cell carcinoma (Li et al., 2011). TAGLN3 encodes a cytoskeleton-associated protein and is reported to possess an association with schizophrenia (Ito et al., 2005). Chromosome 8 shows a selection signal of *myotubularin-related protein 7* (MTMR7) was localized at and associated with the susceptibility of Creutzfeldt-Jakob risk. Three top genes were identified in Chromosome 9, which included *contactin associated protein-like 3B* (CNTNAP3B), *phosphoglucomutase 5 pseudogene 2* (PGM5P2) and *SWI/SNF related, matrix associated, actin-dependent regulator of chromatin, subfamily a, member 2* (SMARCA2). SMARCA2 encodes the protein-controlled coactivator participating in transcriptional activation and vitamin D-coupled transcription regulation. Genetic evidence has shown the association between its genetic polymorphisms and the susceptibility of schizophrenia (Sengupta et al., 2006), Nicolaides-Baraitser syndrome (Van Houdt et al., 2012), lung cancer (Oike et al., 2013). *ADAM metallopeptidase domain 12* (ADAM12) situates in Chromosome 10 and ADAM12 encodes trans-membrane metalloproteinase, which can secret glycoproteins that are involved in cell-cell interaction, fertilisation, and muscle development. We also identified natural selection signatures in *cytochrome P450 family 2 subfamily A member 6* (CYP2A6), *Isthmin 1* (ISM1) and *cytochrome P450 family 2 subfamily D member 6* (CYP2D6).

**Figure 8.**
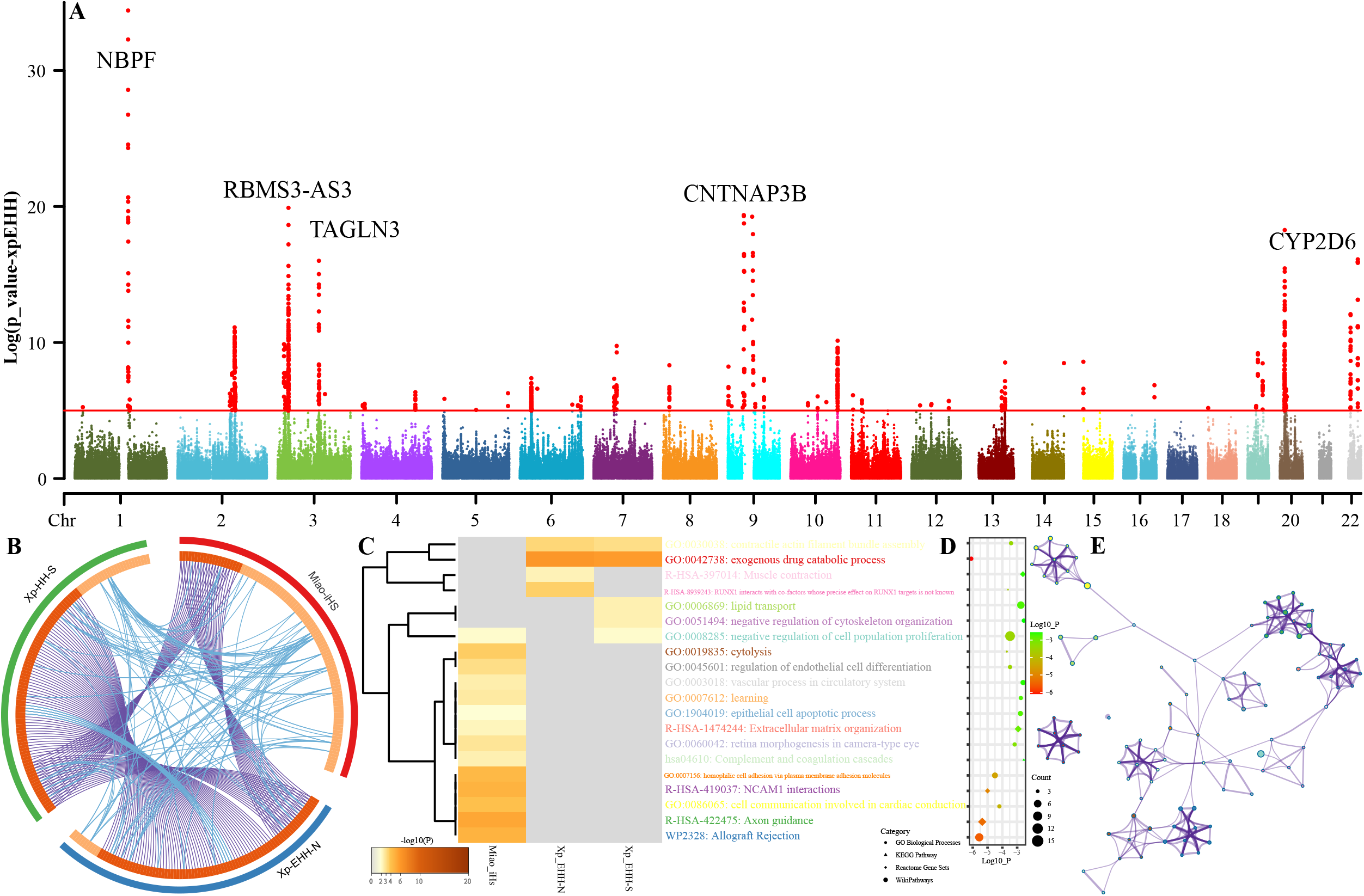
Manhattan showed the natural selection signatures and enrichment analysis. (**A**), P-values of XPEHH in Miao population using northern Han as the reference population. (**B**), Overlap among three gene lists based on gene-level and shared term level, where blue curves link genes that belong to the same enriched ontology term. The inner-circle represents gene lists, where hits are arranged along the arc. Genes that hit multiple lists are colored in dark orange, and genes unique to a list are shown in light orange. (**C**), Heatmap of top-twenty enriched terms across three input gene lists, colored by p-values. (**D**), Top 20 clusters with their representative enriched terms, (**E**), Network of enriched terms colored by cluster-ID.

We further calculated another set of XPEHH scores using southern Han Chinese as the reference population and iHS scores in the SCM populations. To explore the biological functions of all possible natural-selected genes (102 loci in iHS-based, 93 XPEHH_N-based and XPEHH_S-based), we made enrichment analysis based on three sets of identified natural-selection genes. Loci with p-values of XPEHH scores larger than 5 and p-values of iHS larger than 3.3 were used in the enrichment analysis via the Metascape. Overlapping loci observed among three gene candidate lists showed the more common gene candidates inferred from XPEHH and less overlapping loci between XPEHH-based loci and iHS-based loci (**Figure 8B**). Heatmap based on p-values of enrichment pathways (**Figure 8C~E**) showed that all three ways identified the candidate genes associated with metabolic proce ss (GO:0008152), response to stimulus (GO:0050896), cellular process (GO:0009987), regulation of biological process (GO:0050789), biological adhesion (GO:0022610), developmental process (GO:0032502). Results from the iHS also showed other top-level gene ontology biological processes, which included immune system process (GO:0002376), biological regulation (GO:0065007), positive regulation of biological process (GO:0048518), behavior (GO:0007610), signaling (GO:0023052), multicellular organismal process (GO:0032501), locomotion (GO:0040011), negative regulation of biological process (GO: 0048519) and localization (GO:0051179), the detailed enriched terms, pathways and processes enrichment analysis and their networks of top twenty clusters showed in do not reve al the previously reported natural-selected loci associated pigmentation, alcohol metabolism and other common adaptive signals (EDAR et al.) of East Asians (Mao et al., 2021).

## Discussion

### Unique genetic history of HM-speaking populations

Genetic diversity and population history of East Asians have been comprehensively explored and reconstructed in the past twenty years via lower-density genetic markers (Short tandem repeats, SNPs, Indels) and higher-density array-based genome-wide SNPs and whole-genome sequencing data, which advanced our understating of the origin, diversification, migration, admixture and adaptation of Chinese populations (Cao et al., 2020; Chen et al., 2009; Consortium et al., 2009; Wang, C.C. et al., 2021; Xu et al., 2009). As we all know that International Human Genome Organization (HUGO) initiated the broader Human Genome Diversity Project (HGDP) in 1991. HGDP aimed at illuminating the structure of genomes and population genetic relationships among worldwide populations via initial array-based genome-wide SNPs and recent whole-genome sequencing (Bergstrom et al., 2020). A similar work of the CHGDP was publicly reported in 1998 (Cavalli-Sforza, 1998), in which Chu et al. first comprehensively reported genetic relationships and general population stratification based on STR data (Chu et al., 1998). Six years later, Wen et al. illuminated demic diffusion of northern East Asians contributed to the formation of the genetic landscape of modern Han Chinese populations and their sex-biased admixture processes via uniparental markers (Y-chromosome SNPs/STRs and Mitochondrial SNPs) (Wen et al., 2004). The next important step occurred around 2009 and several genetic analyses based on genome-wide SNPs, including mapping Asian genetic diversity reported by HUGO Pan-Asian SNP Consortium, have identified population stratification among linguistic different Asian populations and genetic differentiation between northern and southern Han Chinese populations (Chen et al., 2009; Consortium et al., 2009; Xu et al., 2009). However, these studies had limitations of the lower resolution of used marker panel or limited representative samples from the ethnolinguistic region of China. Recently, large-scale genetic data from the Taiwan Biobank, China Metabolic Analytics Project (ChinaMAP) and other low-coverage sequencing projects (Cao et al., 2020; Chiang et al., 2018; Liu et al., 2018; Lo et al., 2021) have reconstructed fine-scale genetic profiles of the major populations in China and reconstructed a detailed framework of the population evolutionary history. Cao et al. identified seven population clusters along with geographically different administrative divisions (Li et al., 2021), which is consistent with our recently identified differentiated admixture history of geographically different Han Chinese populations possessing major ancestry related to northern East Asians and additional gene influx from neighboring indigenous populations (Guang-Lin He et al., 2021; He et al., 2021; Liu, Y. et al., 2021b; Wang, M. et al., 2021b; Yao et al., 2021). Genetic studies focused on ethnolinguistic Chinese regions further identified different genetic lineages in modern East Asians, TB lineage in Tibetan Plateau, Tungusic lineage in Amur River Basin, AA and AN lineage in South China and Southeast Asia (Siska et al., 2017; Wang, C.C. et al., 2021). Recently ancient genomes also identified differentiated ancestral sources that existed in East Asia since the early Neolithic, including Guangxi, Fujian, Shandong, Tibet, Siberia ancestries (Mao et al., 2021; Wang, C.C. et al., 2021; Wang, T. et al., 2021; Yang et al., 2020). However, many gaps of Southwest Chinese indigenous populations needed to be completed in the Chinese HGDP based anthropological sampling and Trans-Omics of Precision of Medicine of the Chinese population (CPTOPMed). Large-scale genomic data from ethnolinguistic different populations may be provided new insights into the population history and medical utilization in the precision medication for East Asians like the UK10K and TOPMed (Taliun et al., 2021; Wang, Q. et al., 2021).

To comprehensively provide a complete picture of the genetic diversity of China and made comprehensive sampling and sequencing strategies in the next whole-genome sequencing projects, it is necessary to explore the basal genetic background using the small sample size and array genotyping technology. As our part of initial pilot work in CPGDP based on anthropologically-informed sampling, we reported genome-wide SNP data of 55 SCM samples from three geographically diverse populations. Our analysis reveals the key features of the landscape of southwestern HM lineage, including the identified unique HM cline in East Asian-scale PCA and population stratification in regional-scale-PCA, the observed dominant specific ancestry in geographically distant HM people, the estimated strong genetic affinity among HM people via the Fst, outgroup *f_3_*-statistics, *f_4_*-statistics. We further confirmed that stronger genetic affinity within HM people via the sharing patterns of DNA fragments in the IBD, Chromosome painting and FineSTRUCTURE, as well as the attested close clustered pattern in TreeMix - based phylogeny and close phylogenetic relationships between HM people and 500-year-old GaoHuaHua people. Admixture models based on the two-way models further found that the dominant 1500-year-old Guangxi historic ancestry in modern HM people. These observed genetic affinities between HM people from Sichuan, Guizhou, Vietnam and Thailand suggested that all modern HM people possessed a common origin. Combined previous cultural, linguistic, archaeogenetic evidence, the most originated center of modern Hmong-Mine people is the Yungui Plateau in Southeast China. We also found that Miao from Chongqing and HGDP project and She people shared more ancestry with Han Chinese populations, suggested some HM people also obtained much genetic material with southward Han Chinese populations. Compared with historic Guangxi populations (BaBanQinCen and GaoHuaHua), SCM shared more derived ancestry with northern East Asians, suggested that the persistent southward gene flow from northern East Asians influenced the modern genetic profile of HM people. Based on the admixture times dated via GLOBETTROR and ALDER, complex population migration and admixture events occurred in the historic and prehistoric Pro-HM people. Spatiotemporal analysis between modern HM people and their genetic evolutionary relationship with surrounding modern ethnolinguistically diverse populations, as well as the genetic relationship between ancient Yellow River millet farmers and Fujian and Guangxi ancient populations suggested that HM people originated from the crossroad region of Sichuan and Guizhou provinces. Modern HM people may have remained the most representative ancestry of ancient Daxi, Shijiahe people in the middle Yangzi River Basin, which needed to be validated directly via ancient genomes in this region.

### Specific genomic patterns of natural selection signatures

Ethnically different populations undergoing historical differences in the pathogen exposure may remain different patterns of the allele frequency spectrum and extended haplotype homozygosity under natural selection processes. We identified different natural selection candidates (NBPF9, RBMS3-AS3, CNTNAP3B, NBPF10, CYP2D6, TAGLN3, ISM1, RBMS3, KCNJ3, ADAM12, GALNT13, PGM5P2, CYP2A6, MTMR7 and SMARCA2) associated with several different biological functions (metabolic process, response to stimulus, cellular process, regulation of biological processes) in Miao people compared with other East Asians. Denisovan archaic high-altitude adaptive introgression signals were observed in Tibetans (EPAS1 and EGLN1), which is not observed in HM people with obvious natural selection signatures (Yi et al., 2010). More Denisovan archaic adaptive introgression signals related to immune function (TNFAIP3, SAMSN1, CCR10, CD33, DDX60, EPHB2, EVI5, IGLON5, IRF4, JAK1, ROBO2, PELI2, ARHGEF28, BANK1, LRRC8C and LRRC8D and VSIG10L), metabolism (DLEU1, WARS2 and SUMF1) (Choin et al., 2021) were identified in Austronesian and Oceanian populations. But we only observed immune-related Denisovan introgression signals in the DCC gene situated in chromosome 18, which underwent the natural selection evidenced via a higher iHS score (3.5517 in rs17755942, 3.4758 in rs1237775, 3.3540 in rs16920, 3.3299 in rs79301210) in SCM. Choin et al. also reported Neanderthal adaptive introgression genes in Oceanians, including dermatological or pigmentation phenotypes (OCA2, LAMB3, TMEM132D, SLC36A1, KRT80, FANCA and DBNDD1), metabolism (LIPI, ZNF444, TBC1D1, GPBP1, PASK, SVEP1, OSBPL10 and HDLBP), immunity (IL10RA, TIAM1 and PRSS57) and neuronal development (SIPA1L2, TENM3, UNC13C, SEMA3F and MCPH1) (Choin et al., 2021). However, our analysis based on the XPEHH scores only identified one Neanderthal introgression immunity signal (CNTN5) and one pigmentation phenotype signal (PTCH1). CNCN5 harbored high XPEHH scores (>2.1313) ranged from 99577624 to 99616124 in chromosome 11 with the highest values of 4.2829 in rs7111400. Loci situated from 98209156 to 98225683 in PTCH1 in chromosome 9 also possessed higher EPEHH scores in HM people with the highest values in missense mutation rs357564 (4.5412). ALDH2 and ADH1B were reported a strong association with alcohol metabolism (Taliun et al., 2021), however, the highest XPEHH absolute scores in HM people less than 0.5937 for ALDH2 and 1.6013 for ADH1B. Five selection-candidate genes of CTNNA2, LRP1B, CSNK1G3, ASTN2 and NEO15 were evidenced underwent natural selection in Taiwan Han populations (Lo et al., 2021), however, only LRP1B associated with lipid metabolisms were evidenced and replicated in HM people. The observed differentiated patterns of genomic selection process in HM people consistent with their reconstructed unique population history and specific living environments in Southwest China. Thus, further whole-genome sequencing in the CPGDP based on sampling of lager sample size in Southwest China would be provided deep insights of adaptation history of HM people.

## Conclusion

Taken together, we provided genome-wide SNPs data from SCM and directly evidenced their genetic affinity with southmost Thailand and Vietnam Hmong and ancient 500-year-old Guangxi GaoHuaHua people. We identified HM-specific ancestry components spatially distributed ranged from the middle Yangzi River Basin to Southeast Asia, and temporally distributed at least since from 500 years ago. These results provided direct evidence supported a model in which HM-speaking populations originated from ancient Baiyue in the middle Yangzi River Basin and experienced a recent southward migration from Sichuan and Guizhou to Vietnam and Thailand. Additionally, unique patterns of natural-selected signatures in SCM have identified many candidate genes associated with important neural system biological processes and pathways, which not supported the possibility of recent large-scale admixture occurred between HM people and shrouding Han Chinese. If these phenomena occurred, genetic changes can produce shifts in the allele frequency spectrum of pre-existing mutations and trended to showed a consistent pattern of the selected signals.

## Acknowledgments

This work was funded by Supported by the Project funded by China Postdoctoral Science Foundation (2021M691879), Opening project of Medical Imaging Key Laboratory of Sichuan Province (MIKLSP202104), the Science and Technology Program of Guangzhou, China (2019030016), the “Double First Class University Plan” key construction project of Xiamen University (the origin and evolution of East Asian populations and the spread of Chinese civilization), National Natural Science Foundation of China (NSFC 31801040), Nanqiang Outstanding Young Talents Program of Xiamen University (X2123302), the Major project of National Social Science Foundation of China (20&ZD248), the European Research Council (ERC) grant to Dan Xu (ERC-2019-ADG-883700-TRAM). S. Fang and Z. Xu from Information and Network Center of Xiamen University are acknowledged for the help with the high-performance computing. We thank Prof. Wibhu Kutanan in Khon Kaen University, Prof. Mark Stoneking and Dr. Dang Liu in Max Planck Institute for Evolutionary Anthropology for sharing genome - wide SNP data from Vietnam, Thailand, and Laos.

## Data Availability

The Genome-wide variation data have been deposited in the Genome Variation Map (GVM) in Big Data Center, Beijing Institute of Genomics (BIG), Chinese Academy of Science, under accession numbers PRJCA006460 that are publicly accessible at http://bigd.big.ac.cn/gvm/getProjectDetail?project=xxxxxx (available when it published).

